# Quantifying the mechanics of locomotion of the schistosome pathogen with respect to changes in its physical environment

**DOI:** 10.1101/599993

**Authors:** Shun Zhang, Danielle Skinner, Prateek Joshi, Ernesto Criado-Hidalgo, Yi-Ting Yeh, Juan C. Lasheras, Conor R. Caffrey, Juan C. del Alamo

## Abstract

Schistosomiasis is a chronic and morbid disease of poverty affecting approximately 200 million people worldwide. Mature schistosome flatworms wander in the host’s hepatic portal and mesenteric venous system where they encounter a range of blood flow conditions and geometrical confinement. However, the mechanisms that support schistosome locomotion and underlie the pathogen’s adaptation to its physical environment are largely unknown. By combining microfabrication and traction force microscopy, we developed various *in vitro* assays to quantify the mechanics of locomotion of adult male *S. mansoni* in different physiologically relevant conditions. We show that in unconfined settings, the parasite undergoes two-anchor marching mediated by the coordinated action of its oral and ventral suckers. This mode of locomotion is maintained when the worm faces an external flow, to which it responds by adjusting the strength of its suckers. In geometrically confined conditions, *S. mansoni* switches to a different crawling modality by generating retrograde peristaltic waves along its body, a mechanism shared with terrestrial and marine worms. But while the surface of most worms has backward-pointing bristles that rectify peristaltic waves and facilitate forward locomotion, *S mansoni* has isotropically oriented tubercles. This requires tight coordination between muscle contraction and substrate friction but confers *S. mansoni* the ability to reverse its direction of locomotion without turning its body, which is likely advantageous to maneuver in narrow bore vessels. We show that the parasite can also coordinate the action of its suckers with its peristaltic body contractions to increase crawling speed. Throughout this study, we report on a number of biomechanical parameters to quantify the motility of adult schistosomes (e.g. sucker grabbing strength, rate of detachment under flow, peristaltic wave properties and traction stresses). The new series of *in vitro* assays make it possible to quantify key phenotypical aspects of *S. mansoni* motility that could guide the discovery of new drugs to treat schistosomiasis.

## Introduction

Schistosomiasis is a ‘neglected’ infectious disease that affects approximately 200 million people in tropical Africa, South-East Asia and South America (1). The principal species of this flatworm infecting humans are *Schistosoma mansoni, S. haematobium* and *S. japonicum*. Mature parasitic flatworms live in the venous blood system where they mate and produce eggs which eventually exit the body via the feces or urine, depending on species. Many eggs, however, become trapped in internal tissues and organs, inducing chronic inflammation and fibrosis that manifest as pain, malaise and a decreased ability to perform manual or intellectual labor. Given its morbidity, schistosomiasis is a focus of various (inter)national drug delivery campaigns to decrease prevalence and incidence (2,3).

Infection is established via free-swimming larvae (cercariae) that are released from snail intermediate hosts. After penetration of the skin, the cercariae transform to schistosomula (post-infective larvae) and enter the blood circulation, traverse the pulmonary capillaries and establish in the venous system. Worms mature as males or females. The male worm enfolds the female in a specialized gynecophoral canal, and transports the female around the venous system as she deposits her eggs.

Schistosomes possess a muscular system made up of circular, oblique and longitudinal muscles intricately interlaced with a nervous system that utilizes both classical and peptidergic signaling (4-6). This neuromuscular complexity results in a remarkable plasticity in parasite shape and movement (7,8). Further, the parasite possesses anteriorly-positioned oral and ventral suckers which, presumably, help maintain position in the venous system and aid locomotion. Adults of the model schistosome, *S. mansoni*, are found throughout the hepatic portal and mesenteric venous system, which present marked transitions in lumen diameter (*D*) and flow speed (*v*), ranging from *D > 1 cm* and *v ∼ 30 cm/s* within the hepatic portal vein (9) to *D < 0.1 mm* and *v ∼ 1 mm/s* within the mesenteric veins (10). Considering that the body diameter of a mature male worm is approximately 0.5 mm, it is conceivable that crawling and squeezing are employed in the narrow mesenteric veins. In organisms without extremities, crawling and squeezing are typically driven by the exertion of periodic waves of shear stress (*i.e.* traction stress) on the surrounding substrate (11-16). Organisms relying on this locomotor mechanism can control the wave characteristics in order to adapt their migration to diverse properties of their environment, such as surface roughness, adhesiveness and confinement (14,17-20).

The mechanics underlying schistosome locomotion are for the most part unknown. Neither is it known whether and how the flatworm adapts its locomotion to the varying conditions of flow and physical confinement encountered in different parts of the venous system. Specifically, how the pathogen generates traction stress to achieve persistent locomotion is unknown. Nor is it clear whether the parasite employs crawling or squeezing when confined. Moreover, we do not understand whether the pathogen can engage muscle and sucker movement in a coordinated manner. Finally, given that the neuromuscular system of parasitic worms is a rich source of drug targets (5,21), being able to quantify and differentiate the action of experimental chemistries on schistosome locomotion, including relative to the only drug available for treatment, praziquantel, would offer value in the discovery and development of new drugs.

## Methods

### Ethics statement

Maintenance and handling of small vertebrate animals were carried out in accordance with a protocol approved by the Institutional Animal Care and Use Committee (IACUC) of the University of California San Diego. UCSD-IACUC derives its authority for these activities from the United States Public Health Service (PHS) Policy on Humane Care and Use of Laboratory Animals, and the Animal Welfare Act and Regulations (AWAR).

### Preparation of S. *mansoni*

The acquisition, preparation and *in vitro* maintenance of *S. mansoni* have been described (7,22). We employed a NMRI isolate of *S. mansoni* that is cycled between *Biomphalaria glabrata* snails and female Golden Syrian hamsters (Simonsen Laboratories; infected at 4-6 weeks of age) as intermediate and definitive hosts, respectively. Briefly, adult worms were harvested from hamsters 42 days post-infection in RPMI or DMEM, and washed five times prior to maintenance overnight at 37 °C and 5% CO_2_ in Basch medium (23) containing 4% heat-inactivated FBS, 500 μg/ml streptomycin and 500 U/ml penicillin. Worms were maintained *ex vivo* for less than 72 h.

### Polyacrylamide Gel fabrication

Collagen-coated polyacrylamide (PA) gels of 1.5 mm thickness were prepared for traction force microscopy as previously described (24). The gels contained a thin top layer (10 μm) impregnated with 1 μm fluorescent beads (FluoSperes; Molecular Probes) that acted as fiduciary markers of substrate deformation. Gels were fabricated using 5% acrylamide and 0.3% bisacrylamide (Fisher BioReagents), yielding a Young’s modulus of 8.73 kPa. The gels Poisson’s ratio was measured to be 0.46 (25). Gels were activated with sulfo-SANPAH (Thermo Scientific) under UV light and coated with 0.15 mg/ml collagen I (Corning).

### Flow Chamber Experiments

To study *S. mansoni* under conditions representative of their *in vivo* environment, a flow chamber device was fabricated. The device was assembled using 5 laser-cut acrylic sheets (labeled P1 – P5 in Figure 1B), with features tailored to create an inlet that sequentially leads to a reservoir, a rectangular test section that can house a PA gel, another reservoir and the outlet. Silicon grease was applied between each sheet to prevent leaking. The apparatus was fastened by screws.

**Figure 1.**
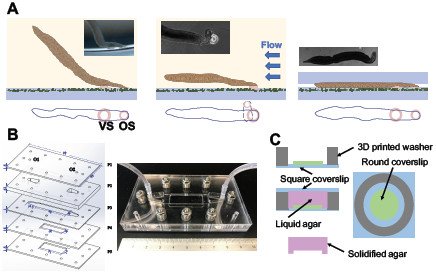
Overview of various experimental designs to model the mechanical environment of *S. mansoni*. (**A**) Schematics of a free adhesive (sucker-based) locomotion (left), a worm adhering against flow (center), and a worm moving under confinement (right), with corresponding bright-field images (left, lateral view; center and right, dorsal views) captured during our experiments: VS and OS refer to the ventral and oral suckers, respectively. (**B**) Flow chamber design. The image to the left illustrates the device’s layered design with the important dimensions indicated in mm. O1 and O2 are inlet and outlet orifices. The image to the right is a photograph of the assembled device. (**C**) Fabrication scheme of the agar cap system used to generate controlled confinement. A 3D printed PLA washer (grey) and a round coverslip (green) are glued onto a square coverslip (blue). An agar solution (purple) is poured and covered with a square coverslip to flatten its surface while the solution solidifies. Varying the number of round coverslips controls the gap produced in the agar gel.

Prior to each experiment, the device was filled with culture medium. Subsequently, a 1 ml pipette was used to gently introduce 8-10 adult male worms into the device from the inlet. The inlet was then connected to a 60-ml syringe (BD) filled with culture medium, which was gently pushed to drive the worms into the test section. Then the device was placed onto the stage of an inverted microscope for image acquisition with the syringe installed on a NE-4000 syringe pump (Pump Systems Inc).

The flow rates in our experiments were chosen to cover the physiological wall shear stress encountered in the human venous system (from the vena cava to the venules) (26). A flow rate of *Q =* 2.24 ml/min was applied after *S. mansoni* adhered to the PA substrate. Then the flow rate was increased in a stepwise manner to *Q =* 11.2, 56.1 and 100 ml/min with each flow step lasting for at least 10 seconds. At each flow step, worms that were able to adhere to the PA substrate >5 seconds were classified as adhering worms. The recording was stopped upon worm detachment or after reaching the maximum flow rate. We followed the Boussinesq velocity profile in a rectangular channel (27) to calculate flow properties from *Q* and chamber geometry, including Reynolds number, maximum flow velocity, wall shear stress and pressure gradient (Table 1). Culture medium viscosity was measured as *μ = 7.1****×****10*^*-4*^ Pa s at 37° C using a rheometer (*micro*VISC).

**Table 1.**
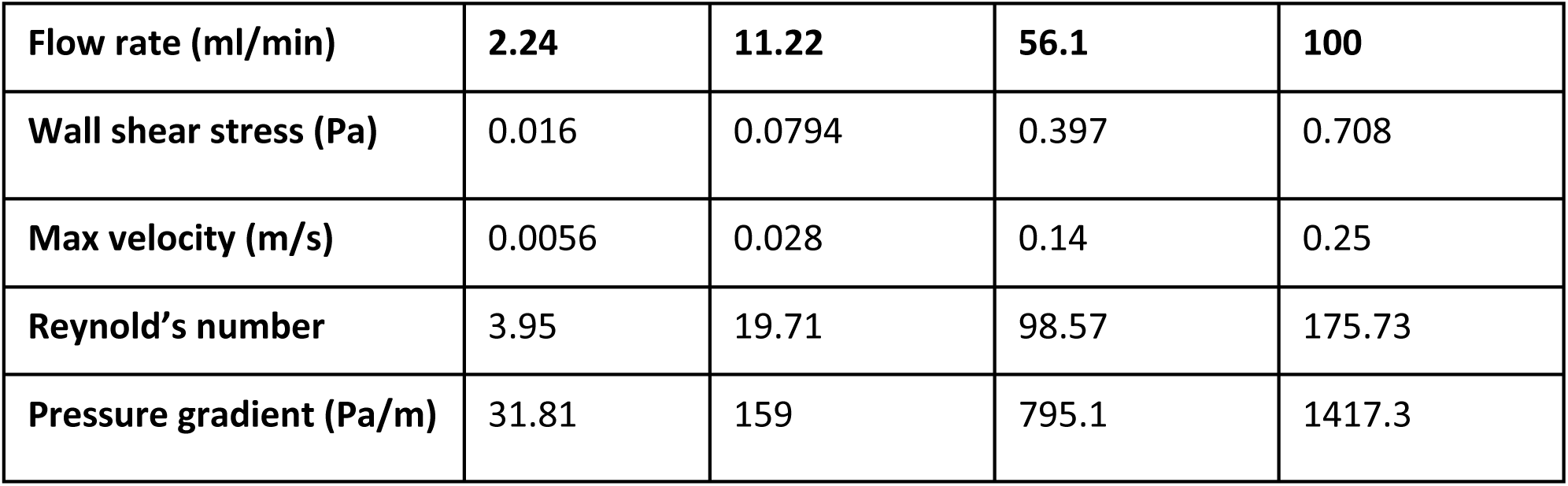
Calculated relevant flow properties with flow rate *Q =* 2.24, 11.2, 56.1 and 100 ml/min.

Among all of the vessels encountered by *S. mansoni*, the portal vein has the highest flow velocity. A reference wall shear stress value in the hepatic portal vein (*WSS*_*hpv*_) (9) was calculated by assuming a Poiseuille profile with peak velocity of 30 cm/s, a vessel diameter of 1 cm and a blood viscosity of μ_blood_ = 2.73 x 10^−3^ Pa s at 37° C, resulting in a value of 0.33 Pa. This value falls within the range of wall shear stresses induced in our device.

### Experiments in a confined environment

To mimic the physically restrictive environment encountered by *S. mansoni* in narrow veins, we sandwiched the worms between the PA gel used for traction force microscopy and an agar cap (0.8% agarose, 18 mm diameter and 1.5 mm thickness), as in our previous work (17,18). To generate varying levels of confinement, the worms were sandwiched in a gap of either 300 or 100 μm (Figure 1C). Considering that the thickness of an adult male *S. mansoni* is ∼ 500 μm, these gaps confined the worm gently or severely, while still permitting worm motion. Caps were soaked in culture medium for 30 minutes before experiments, then placed on top of the PA gels immediately after transferring the worms.

### Microscopy

A Leica DMI 6000B inverted microscope controlled by Micro-Manager software was used for image acquisition (28). All experiments were performed at 37 °C. Different acquisition protocols were followed as listed in Table 2. The experiments performed under no confinement (including the flow chamber experiments) were imaged under 5X at 1 frame per second (fps) in both the bright and fluorescent fields. This field of view was sufficient to capture the oral and ventral suckers. Under confinement, we imaged at 2X in order to image the whole length of the body. Also, we acquired time-lapse image sequences at 10 and 100 fps in bright field only.

**Table 2.** Acquisition protocols followed in this study.

| **Experimental Design** | **Channel** | **Frame rate (fps)** | **Magnification** | **Purpose** |
| --- | --- | --- | --- | --- |
| **No constrain, no flow** | BF & FLUO | 1 | 5X | Movement, traction stress under sucker |
| **No constrain, with flow** | BF & FLUO | 1 | 5X | Traction stress under sucker |
| **Constrain, no flow** | BF & FLUO | 1 | 2X | Movement and body wave, traction stress under body and sucker |
| **Constrain, no flow** | BF | 10 | 2X | Higher temporal resolution of body wave |
| **Constrain, no flow** | BF | 100 | 5X | Velocity of material points on the body |

### Traction Force Microscopy

The horizontal, *i.e*., in-plane, deformation of the top surface of the PA gel was measured by tracking the displacement of the fluorescent beads as reported previously (24). Each instantaneous image was correlated with a reference image recorded after the worms moved out of the field of view. Using these measurements as boundary conditions and assuming zero vertical deformation, we solved the elastostatic equation for the PA gel as previously described (24,29). This solution provided the horizontal traction stress vector *τ =* (*τ*_*xz*_, *τ*_*yz*_) on the top surface of the PA gel. The spatial resolution (i.e. the distance between consecutive measurement points) was 20 μm and 50 μm under 5X and 2X, respectively.

In our experiments, the vertical deformation induced by the worms was not negligible. However, acquiring 3D image stacks at a frame rate high enough to capture the dynamics of worm locomotion (∼1 fps) was unfeasible in our setup. Thus, we chose the substrate’s Poisson’s ratio and its thickness relative to worm size so that the horizontal and vertical components of the traction stress were decoupled, based on existing theoretical predictions (24). Consequently, our 2D TFM experiments can still be used to accurately measure in-plane traction stress.

### Kymograph Representation

We generated kymographs to analyze the spatio-temporal dynamics of *S. mansoni* locomotion. At each instant of time (*t*), the centerline of the worm’s body was determined from bright field images. Then, the distance along the centerline from the posterior tip of the worm was used as an independent variable together with time to create 2D representations by stacking together data from different time instants. In particular, we plotted spatiotemporal distributions of worm width *w*(*s,t*) and longitudinal traction stress *τ*_*s*_ (*S,t*,) ***= τ*_*w*_** (*S,t*,) · ***e*_*s*_** (*S,t*,), where ***τ*_*w*_** represents the local average of the traction stress across the worm’s body width and ***e*_*s*_** is a unit vector parallel to the body’s centerline.

## Results

We studied the mechanics of locomotion of *S. mansoni* in three different physiological environments (Figure 1A): (i) no confinement and no flow, (ii) no confinement under directional flow and (iii) geometrical confinement.

### The suckers mediate schistosome adhesion and locomotion in unconfined environments

When placed on a flat surface in the absence of confinement, adult males alternated resting phases, during which they adhered with the oral and/or ventral sucker, with “marching” bouts that involved the coordinated adhesion of both suckers and contractility of the neck region between them (Figure 2). Both suckers generated various patterns of traction stress on their substrate, namely contraction, expansion and rotation. Most frequently, we recorded an inward contractile pattern (Figure 2A, D), consistent with a suction pressure being generated inside the sucker. This pattern was also observed under flow and confinement, as shown below. Outward stress patterns also appeared occasionally (Figure 2B, E) but these events were transient and did not occur under flow. The data suggest that the suckers of S. *mansoni* are sessile and generate pressure differences by expanding and contracting a suction cup, in contrast to the piston-like stalk suckers found in some cephalopods, which can create larger pressure differences (30).

**Figure 2.**
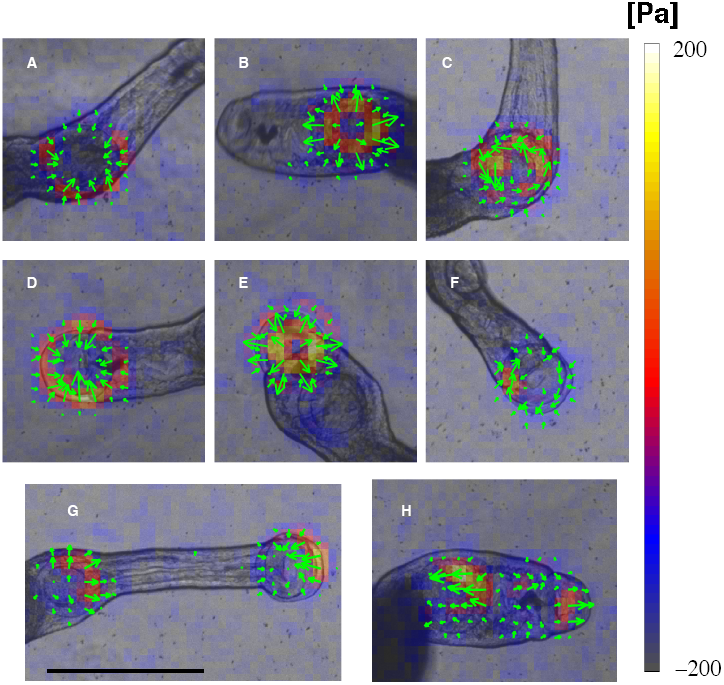
Instantaneous traction stress patterns applied by S. *mansoni* suckers in an unconfined environment. (**A, B, C**) Inward, outward and rotational patterns induced by the ventral sucker alone. (**D, E, F**) Inward, outward and rotational patterns induced by the oral sucker alone. (**G**) Contractile traction stress pattern applied by coordination of the oral and ventral suckers. (**H**) Expanding traction stress pattern applied by coordination of the oral and ventral suckers. The arrows and color map indicate the direction and magnitude of the traction stress vector, and have been superimposed on the bright field images. Scale bar = 1 mm.

Rotational traction stress patterns (Figure 2C,F) were observed under the suckers coinciding with changes in the worm’s direction of motion or with pivoting of the body. To understand whether the suckers actively generated torque or whether the observed rotational patterns were a passive reaction to the hydrodynamic drag acting on the worm’s body, we performed experiments with “body-less” worms that had been cut just posterior to the ventral sucker. The suckers of these fragments still generated contractile and expansive traction stress patterns in the suckers but not rotational ones, implying that suckers are unable to actively generate torque.

When marching forward, the neck elongated supported by the ventral sucker until the oral sucker adhered to the substrate. Then, the ventral sucker detached and the neck contracted, pulling the entire worm forward towards the oral sucker, which, in turn, detached after the ventral sucker re-attached (Supplementary Videos 1-4). The reverse sequence of actions was observed for backward marching (Supplementary Video 5). In many instances, while marching, the neck elongated or contracted before one of the suckers had detached from the substrate, which allowed us to measure a pattern of contractile (Figure 2G) or expansive (Figure 2H) traction stress sustained across the whole neck. These forces might drive the detachment of the suckers, suggesting that they may sense mechanical forces to regulate locomotion.

### Sucker adhesion strength increases in response to increasing flow stress

To study the adhesion and motility of *S. mansoni* under physiological flow conditions, we performed experiments in a custom-built device (Figure 1A2). We quantified the number of worms that remained attached to the substrate as the flow rate (*Q*) and wall shear stress (*WSS*) were progressively increased (Figure 3A). The percentage of worms able to withstand the hydrodynamic drag without detaching decreased sharply with increasing *Q*; whereas >80% of the worms remained attached at the lowest *Q* = 2.24 ml/min, only ∼20% of the worms remained attached at the highest *Q* = 100 ml/min. Furthermore, the worms transitioned from using either the oral or the ventral sucker alone to grab the substrate under no flow, to preferentially using the stronger ventral sucker, and eventually both suckers as *Q* was increased (Figure 3B). Because the effect of increasing *Q* on worm detachment was reminiscent of the dose-response curve of a drug effect, we fitted our data using the Hill dose-response equation (31,32) and found that the 50% detachment flow rate was *Q*_*50*_ *= 26.6 ml/min*. This flow rate is equivalent to a wall shear stress *WSS*_*50*_ *= 0.19 Pa*, which is ∼40% lower than the typical value of the *WSS* in the hepatic portal vein (*WSS*_*hpv*_ *= 0.33 Pa*, see Methods Section).

**Figure 3.**
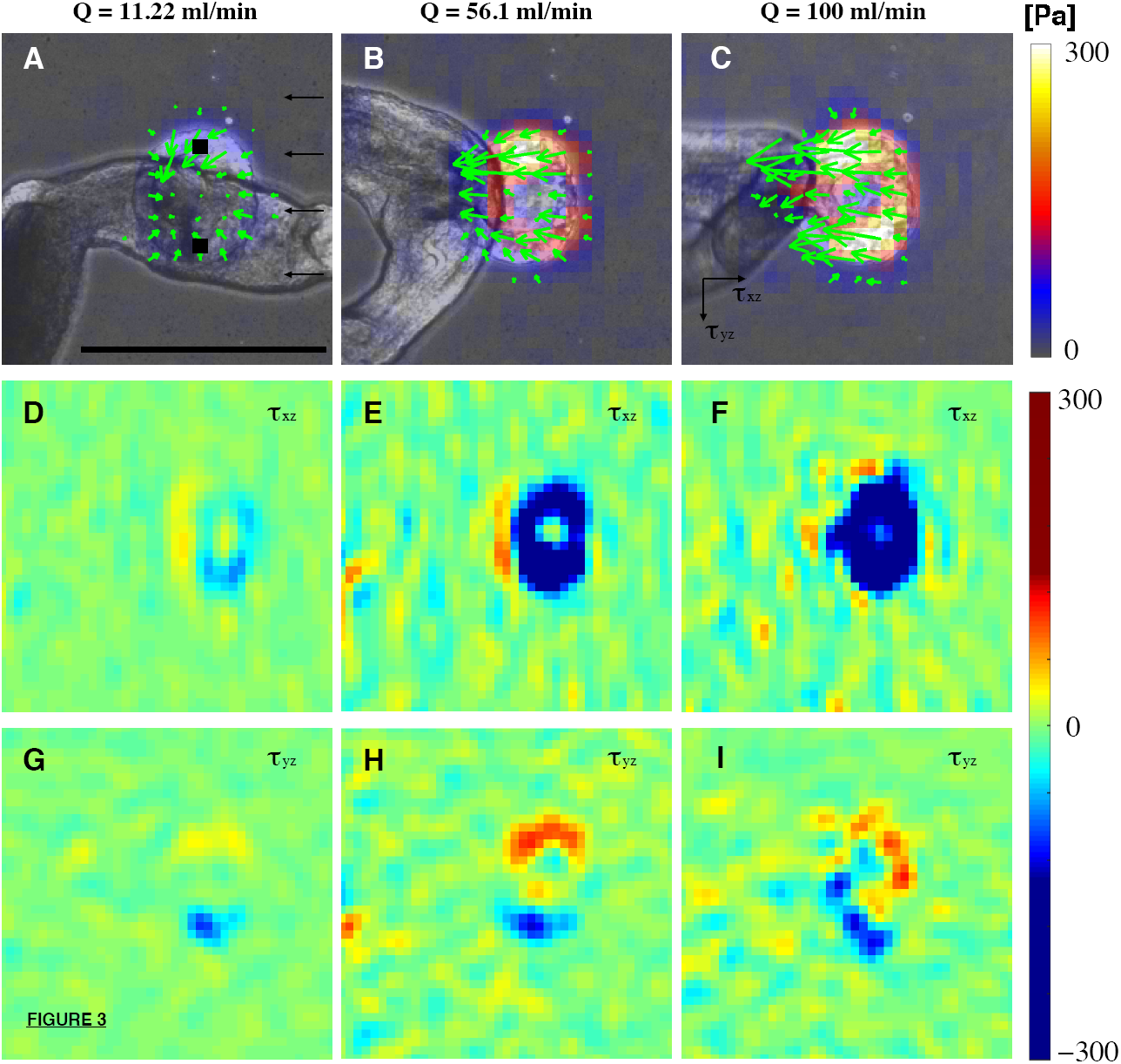
Substrate adherence of *S. mansoni* under flow. (**A**) Percentage of adherent S. *mansoni* as a function of flow rate and corresponding wall shear stress (average +/-standard error, n=4 worm batches with 11, 20, 5 and 5 worms). A dose-response curve based on fitting to the Hill Equation is plotted in black. The blue square indicates the 50% detachment flow rate based on the model. (**B**) Distribution of sucker usage as a function of flow rate and corresponding wall shear stress (Blue – oral sucker alone; green –ventral sucker alone; red –both suckers. Data are average +/-standard error, n=41 worms).

Our traction stress measurements (Figure 4) were consistent with the observed response in attachment. At low *Q*, the ventral sucker exerted a contractile pattern of traction stress, as indicated by the dipolar structure of both the flow (*τ*_*xz*_) and cross-flow (*τ*_*yz*_) components of the stress (Figure 4A, D, G). This pattern was similar to that observed in the flow-free condition (Figure 2A). As *Q* increased, so did the hydrodynamic drag experienced by the worm, which was balanced by the traction stress under the ventral sucker. Consequently, *τ*_*xz*_ increased in magnitude and lost its dipolar structure, becoming unidirectional (Figure 4D-F).

**Figure 4.**
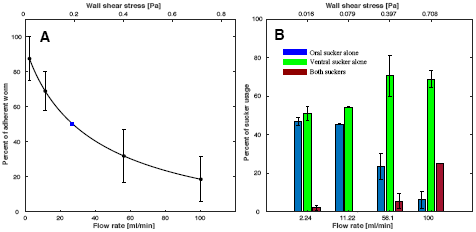
Instantaneous traction stress patterns applied by S. *mansoni* anchoring against flow. (**A, B, C**) Traction stress patterns for different flow rates. The green arrows and color map indicate the direction and magnitude of the traction stress vector. (**D, E, F**) Traction stresses in the flow direction (*τ*_*xz*_) for different flow rates. (**G, H, I**) Traction stresses perpendicular to the flow direction (*τ*_*yz*_) for different flow rates. (**A, D, G**) Flow rate = 11.22 ml/min, (**B, E, H**) 56.1 ml/min and (**C, F, I**) 100 ml/min. The black squares in **A** indicate positions where grabbing strength is calculated from traction stress. The black arrows in **A** indicate the direction of flow. The black arrows in **C** indicate the direction of traction stress. Scale bar = 500 μm.

According to these data, there should be a maximum level of traction stress *τ*_*max*_ that the ventral suckers of *S. mansoni* can withstand before losing substrate adherence. By applying force balance for the highest flow rate condition, we estimated that

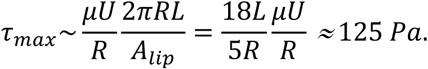

This estimation assumed that (i) the flow around the worm is dominated by viscous stress (*i.e*., steady flow with negligible inertia), (ii) the worm is cylindrical with a length *L ∼ 10*^*-2*^ *m* and a radius *R ∼ 0.25 x 10*^*-3*^ *m*, (iii) R is negligible with respect to the channel height H, i.e. *R<<H*, (iv) *μ ∼ 10*^*-3*^ *Pa s* is the viscosity of the medium, (v) *U ∼ 0.25 m/s* is the centerline velocity in the flow chamber, (vi) 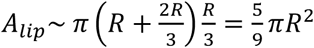 is the area of the sucker lip in contact with the substrate, and (vi) the lip has a width equal to *R/3* (Figure 1A). Our estimate reasonably agrees with the traction stress values measured in the flow chamber for the highest *Q* (Figure 4F). Using similar arguments, the estimate of *τ*_*max,50*_ corresponding to *Q*_*50*_ *= 26.6 ml/min* is *τ*_*max,50*_ *∼ 35 Pa*. It should be noted that our assumption *R<<H* holds only approximately in our *in vitro* experimental setup where *R/H ∼1/2*, and likely breaks down for narrow-bore veins *in vivo*. The effect of confinement (R∼H) may cause the detachment stresses to be lower than our estimation (33,34).

Interestingly, the cross-flow traction stress kept a dipole pattern that balanced out to zero as *Q* was increased but it also increased in magnitude (Figure 4G-I; the slight departures from symmetry were most likely caused by the sideways deflection of the neck region). This result implies that *S. mansoni* senses hydrodynamic forces and responds to them by grabbing the substrate with increasing strength in order to remain attached to it. To test this hypothesis, we plotted the grabbing strength of each worm vs. *Q* normalized by its corresponding value for *Q = 0* (Figure 5). We quantified grabbing strength as the average magnitude of *τ*_*yz*_ at two diametrically opposed regions under the adhering ventral sucker in the cross-flow axis (black regions in Figure 4A). We then used our estimation of *Q*_*50*_ to categorize each worm as adherent if it remained attached for Q > *Q*_*50*_ or non-adherent otherwise. In agreement with our hypothesis, adherent worms were able to increase their grabbing strength gradually in response to increasing *Q*, whereas non-adherent worms did not adjust their grabbing strength.

**Figure 5.**
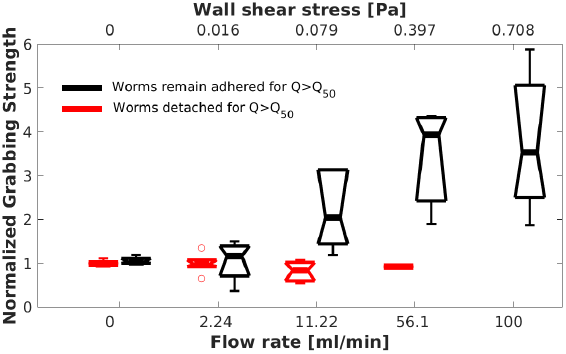
Adhesive strength of *S. mansoni* subjected to stepwise increases in flow rate. The adhesive strength is calculated from the traction stresses perpendicular to the flow direction and normalized to the value at a zero flow rate. The black boxplots correspond to worms that remain attached to the substrate for flow rates Q >*Q*_*50*_ (n=7 worms). The red boxplots correspond to worms that detached from the substrate for Q >*Q*_*50*_ (n=6 worms).

### Locomotion of *S. mansoni* under varying degrees of confinement

To study the motility of *S. mansoni* in conditions representative of the narrower bores of mesenteric veins, we sandwiched worms between a PA substrate and agar caps that created varying levels of confinement. The traction stress exerted by the worms varied notably with the level of confinement. Under no confinement, S. *mansoni* employed its suckers to move and no measureable traction stress was recorded along the main body (Figure 6A-D). Under the gentle confinement generated when the gap between the PA gel and the agar cap was slightly narrower (300 μm) than the worm’s body diameter (∼500 μm), *S. mansoni* maintained the same two-anchor mechanism used on free surfaces. However, the traction stress applied by the suckers increased by over an order of magnitude to account for the friction created between the body and the substrate (Figure 6E-H).

**Figure 6.**
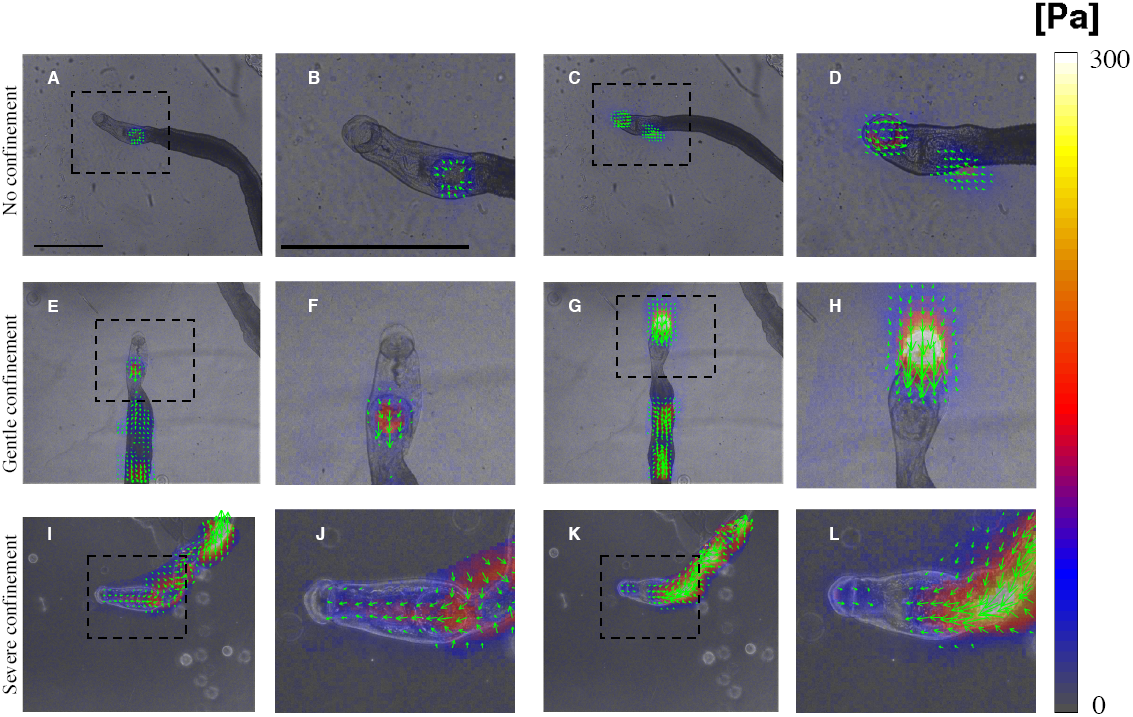
Instantaneous traction stress patterns exerted by *S. mansoni* under varying levels of confinement. (**A-D**) No confinement. (**E-H**) Gentle confinement (300 μm gap between substrate and agar cap). (**I-L**) Severe confinement (100 μm gap). (**A, E, I**) Traction stress exerted while extending the oral sucker forward. (**C, G, K**) Traction stress exerted while contracting the neck region between the oral and ventral suckers. Panels **B, D, F, H, J** and **L** are enlarged views of the boxed areas in panels **A, C, E, G, I** and **K**, respectively. The arrows and color map indicate the direction and magnitude of the traction stress vector, and have been superimposed on the bright field images. Scale bar = 1 mm.

Strikingly, S. *mansoni* switched to a completely different locomotion strategy when crawling under the severe confinement generated when the gap between the PA gel and the agar cap was decreased to 100 μm. Specifically, we observed metachronal waves of body width associated with traveling waves of strong traction stress along the body of the worm (Figure 6G-L). These waves exhibited a rich variety of spatio-temporal patterns leading to a wide range of locomotion behaviors (Supplementary Videos 6-8). We focused on persistent forward locomotion which was always achieved by peristaltic waves of muscle contraction.

Instantaneous worm snapshots (Figure 7A) and the corresponding kymographs of body width (Figure 7B) revealed that 2-3 peristaltic waves were present along the worm’s body at any given time. The waves always originated immediately behind the ventral sucker (green dot in Figure 7A), and traveled backwards with a wave speed *c ≈ 1 mm/s*. The velocities of material points on the surface of the worm were measured by tracking their associated texture in bright-field images and mapped in Figure 7C. Specifically, we found that there are many dark spots distributed on the worm’s body, which can be used as fiduciary makers to measure the velocities and strains of worm body, similar to Lai *et al* (14). The instantaneous longitudinal strain of the worm’s body was estimated from the measured velocity field by assuming a traveling wave profile *u = U*(*x -ct*,), where *u* is the longitudinal displacement of the body, x is the Eulerian distance along the body axis, and *c* is the wave propagation speed. Consequently, Figure 7C also maps the strain 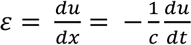. These data indicate that peristaltic muscle contractions locally shortened the worm’s body longitudinally, which in turn became wider to conserve hydroskeletal volume (red bands in kymography). Conversely, the interwave regions were longitudinally stretched and radially narrowed. The data also suggest that wave regions experienced very slow (∼10 μm/s) backward displacement and were effectively anchored to the substrate, whereas interwave regions experienced fast (∼100 μm/s) forward displacement.

**Figure 7.**
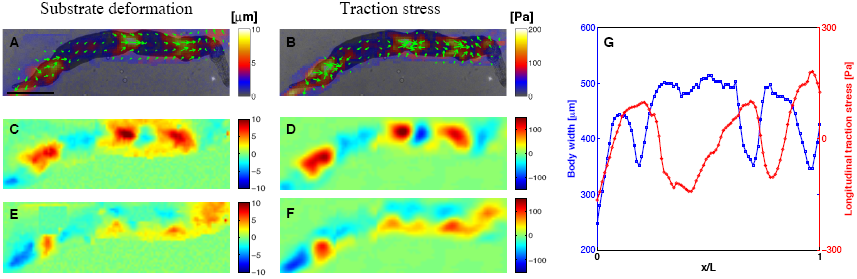
Kinematics of *S. mansoni* movement under severe confinement. (**A**) Instantaneous body width of a worm (represented in color along the worm’s centerline) in a 100 μm gap showing three contraction waves. The green and magenta dots indicate the starting (just behind the ventral sucker) and final (500 μm from the posterior tip) positions of the body width measurement. (**B**) Kymograph of body width represented as a function of time and normalized position along the centerline (1=anterior end and 0=posterior end). (**C**) Instantaneous map of body movement velocities. The arrows represent the direction and magnitude of the velocity vector. The color maps indicate the velocity and estimated strain (ε) in the direction along the worm’s centerline. The arrows and color map have been superimposed on the bright field images. Scale bar = 1 mm.

Overall, our findings indicate that *S. mansoni* exploits friction to generate thrust forces at the wave regions, and that these forces balance the resistance created by the forward displacement of the interwave body regions. To confirm this idea, we measured the deformations and traction stress during worm locomotion and mapped them together with bright field images of the worm body (Figure 8A, B). As expected, these maps showed that the longitudinal substrate deformations and traction stress were propulsive, *i.e*. pointed backwards, in the wave regions, whereas they were resistive in the interwave regions (Figure 8C, D). The data also confirmed that the interwave regions co-localized with regions of radial narrowing (Figure 8E, F, G). Kymographs of jointly acquired body width and longitudinal traction stress (Figure 9) confirm that both quantities exhibited retrograde traveling waves with the same period and wave speed, and with opposite phases. Of note, the worms could move backwards by reversing the direction of wave propagation (Supplementary Video 8).

**Figure 8.**
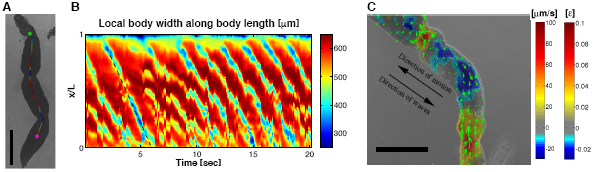
Dynamics of *S. mansoni* movement under severe confinement. (**A**) Instantaneous substrate deformation patterns exerted by a moving worm in a 100 μm gap. (**B**) Instantaneous traction stress patterns exerted by a moving worm. (**C, E**) Maps of instantaneous substrate deformation in the directions longitudinal (**C**) and transverse (**E**) to worm body motion. (**D, F**) Maps of instantaneous traction stress in the directions longitudinal (**D**) and transverse (**F**) to worm body motion. In panels (**A**) and (**B**), the arrows and color maps represent the direction and magnitude of the deformation and traction stress vectors, and have been superimposed on bright field images of the worm. Scale bar = 1 mm. (**G**) Simultaneous measurements of worm body width (blue) and longitudinal traction stress (red) along the worm’s centerline (0 = posterior end, 1 = anterior end).

**Figure 9.**
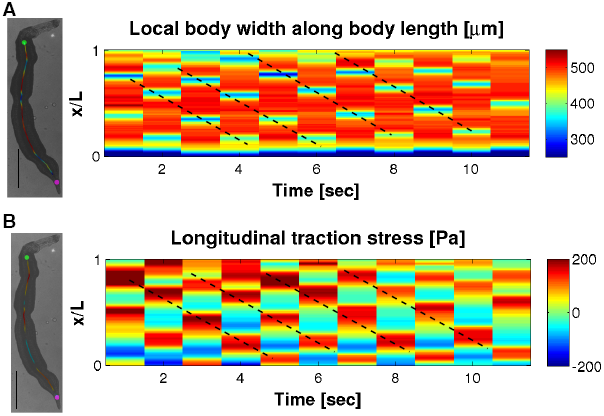
Peristaltic waves of body contractions and propulsive stresses. (**A**) Kymograph of *S. mansoni* body width moving under severe confinement (100 μm gap). (**B**) Kymograph of longitudinal traction stress recorded simultaneously with body width for the same worm. The dashed lines in both kymographs correspond to peristaltic waves of body width and longitudinal traction stress propagating backward. The worm images to the left show instantaneous body width and traction stress measurement along the worm’s centerline.

### Sucker action and peristaltic body contractions can be coordinated during locomotion under confinement

During locomotion in confined environments, about 75% (14 out of 19) of the worms employed peristaltic body waves and not their suckers. The other 25% employed their suckers in addition to body waves (Supplementary Videos 10, 11). To understand whether these two independent mechanisms could be coordinated, we plotted vs time the length of the neck region and the worm’s body width immediately posterior to the ventral sucker (Figure 10A). The data indicate that, in worms using both their suckers and peristaltic waves, the waves were initiated precisely when the ventral sucker completed its forward step and the neck region reached maximum contraction (Figure 10B and Supplementary Video 9). Consequently, a continuous wave of forward body displacement was generated along the whole length of the worm in this locomotion regime (Figure 10C).

**Figure 10.**
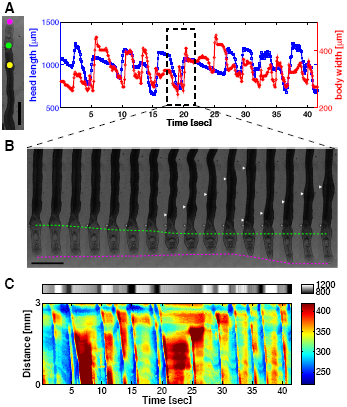
Coordination between peristaltic body contractions and sucker-mediated propulsion in *S. mansoni* under confinement. (**A**) Head length (area between the oral and ventral sucker) of a worm as a function of time (blue) together with body width at a longitudinal position near the head (red) under confinement (100 μm gap). The magenta and green points in the bright field image to the left mark the anterior and posterior ends of the head region, whereas the yellow point marks the location at which body width was plotted. Scale bar = 1 mm. (**B**) Montage of bright field images showing how peristaltic waves of body contraction are initiated whenever the head region reaches its minimum length. White arrowheads mark the position of body wave, and the green and magenta dashed lines indicate the positions of ventral and oral suckers, respectively. (**C**) Composite kymograph of head length (top, grey color map) and body width (bottom, rainbow color map). The data show multiple instances when peristaltic body contraction waves (blue ridges) firing at the point when the head length is shortest (dark grey).

Worms with sucker-peristaltic wave coordination migrated significantly faster than those employing peristaltic waves alone (median speed 80.6 μm/s vs. 25.7 μm/s; Mann-Whitney test *p*-value < 0.01; Figure 11A). This difference may be explained by considering that worms advance on average a fixed stride per wave period (the slope of the regression lines in Figure 11B). Then, engaging the suckers in the motion cycle results in a longer stride (174 μm vs 95 μm, Figure 11B). In an attempt to find additional differences between the two locomotion modalities, we compared their wave frequency, wave speed and worm body length. Because we found a trend for wave frequency to be higher in worms with sucker-peristaltic wave coordination, we compared wave speed and body length in a subgroup of experiments in which the wave frequency was more than 0.3 Hz. However, we did not find significant differences in these variables (Figure 11C, D, E).

**Figure 11.**
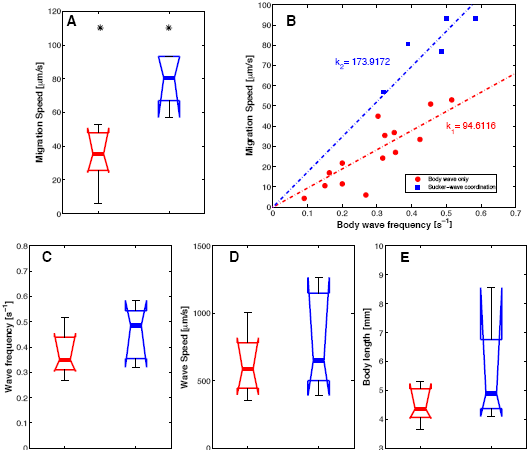
Coordination between peristaltic body contractions and sucker-mediated propulsion modulates speed of movement under severe confinement. (**A**) Box plots of *S. mansoni* worm speeds (in a 100 μm gap) with a wave frequency of more than 0.3 Hz using exclusively peristaltic waves of body contractions (red, n=8) or by coordinating sucker motion with peristaltic body waves (blue, n=5). Asterisks denote statistically significant differences between medians (Mann-Whitney test, *p*-value < 0.01). (**B**) Scatter plot of movement speed versus frequency of peristaltic waves. Red circles - peristaltic waves only; blue squares – coordination between the suckers and peristaltic waves. Linear least square fits through the origin are applied to each data set. (**C-E**) Box plots of motility parameters for worms with wave frequency more than 0.3 Hz migrating exclusively by peristaltic waves of body contractions (red, n=8) or by sucker - peristaltic wave coordination (blue, n = 5); (**C**) average wave frequency, (**D**) average wave speed and (**E**) average body length.

## Discussion

During its life cycle, the schistosome parasite employs a diversity of modalities to migrate within and between its hosts. Eggs secrete hydrolytic enzymes and influence the host’s immune response to migrate through tissues and egress into the gut or bladder lumen (35-38). The free-swimming miracidium uses cilia to move through water. Penetrance of snails is achieved via both mechanical and hydrolytic mechanisms (4,39,40). The emerging infective larvae (cercariae) possess a forked tail to remain suspended in the water column (4,41) until contact with a definitive vertebrate host, including humans. Cercariae then penetrate the skin employing both mechanical and hydrolytic methods (4,42-44). Adult schistosomes are known to wander extensively in the mesenteric system (4,45), whereby the male ‘chauffeurs’ the female around as she performs her primary task of laying eggs. However, the mechanics of locomotion of adult male worms have not been formally evaluated, much less quantified.

### The versatile mechanical modalities of locomotion in *S. mansoni*

This study demonstrates that the adult male *S. mansoni* flatworm engages various mechanical principles to generate remarkable locomotor versatility under different environmental conditions.

#### Two-anchor marching under external flows

The primary conveyance method in unconstrained environments is two-anchor marching mediated by the worm’s suckers, both under flow and no-flow conditions (Figure 2 and Supplementary Videos 1-4). By alternating the attachment and release of its oral and ventral suckers, *S. mansoni* can remain securely anchored to the substrate while marching, thus avoiding being washed away by external flow. With increasing hydrodynamic resistance, *S. mansoni* transitions from using either sucker to adhere to the substrate, to using the exclusively the ventral sucker or both suckers (Figure 3), implying that the main function of the ventral sucker is to hold the pathogen in place. The oral sucker is smaller in size and has a more versatile motor function. It is also a mouth through which the parasite feeds on blood materials (4). Our traction stress measurements in flow chamber confirmed the fundamental importance of the ventral sucker in withstanding hemodynamic forces (Figure 4). Furthermore, our experiments revealed that *S. mansoni* senses these drag forces and respond to them by increasing the strength with which the sucker grips the substrate (Figure 5). The ability to sense drag is a novel finding and consistent with the complexity of the pathogen’s neuromuscular system.

#### Reversible peristaltic locomotion as an adaptation to squeeze through constricted environments

Under physical confinement, we observed that *S. mansoni* switches from sucker-based marching to propulsive peristaltic waves (Figure 6 and Supplementary Videos 6-8). This is a useful adaptation as the parasite seeks to squeeze through the mesenteric venous system in order to deposit eggs as close as possible to the intestine and thereby facilitate parasite transmission. During forward locomotion, waves of muscle contraction are typically fired posteriorly from the ventral sucker and propagate along the main body of the parasite (Figure 7B and Supplementary Video 6). Conversely, for reverse locomotion, the waves originate at the rear of the worm and travel toward the ventral sucker (Supplementary Video 8). In both cases, we found that the peristaltic waves propagate in the direction opposite to whole worm locomotion, *i.e*., are retrograde waves (46). Our simultaneous measurements of body width, motion velocity and longitudinal strain (Figure 7C) indicate that the worm effectively anchors to the substrate in the peristaltic wave regions, which slip backward very slowly, and are subjected to longitudinal contraction and radial swelling. These wave regions exert the propulsive forces needed to balance the resistance force caused by the forward displacement of the interwave regions (Figures 8-9), where elastic recoil from longitudinal stretch likely drives fast forward displacement of the body.

#### Increased locomotion involving synchronized marching and squeezing

Our experiments revealed that the adult *S. mansoni* male worm possesses considerable locomotor plasticity. Particularly, the parasite can not only adjust its locomotion to its physical environment but can also simultaneously combine different gaits and modalities. For instance, we observed that worms combine forward and backward propagation of peristaltic waves in the anterior and posterior parts of their body to achieve sideways locomotion (see Supplementary Video 7). Moreover, we found that worms can synchronize sucker-mediated locomotion and peristaltic body waves to achieve significantly faster locomotion under confinement (Figures 10-11 and Supplementary Videos 9-11). Mechanically, this “blended” locomotion mode is efficient because 1) it allows the suckers to contribute to locomotion in a restrictive physical environment while still exerting low traction stresses, 2) it abolishes the resistance force created by the otherwise passive head region of the worm, and 3) the synchronous succession of head extension and firing of peristaltic waves leads to an increased stride length (Figure 11B). Our recent studies reveal that *S. mansoni* has a unique brain among flatworms that comprises two pairs of cerebral ganglia, one pair forward of the ventral sucker and the other aft of the oral sucker (5). These data pose a fundamental question for brain function involved in controlling the performance of and interplay between body wave- and sucker-based locomotion.

### Coordination between peristaltic contraction and substrate friction

Our measurements of traction stress and worm body motion (Figures 6-9) suggest that both the propulsive and resistive forces that govern the locomotion of *S. mansoni* are transmitted onto the substrate by friction. These results imply that the parasite can regulate the coefficient of friction with its surroundings in coordination with muscular contraction. Otherwise, the global balance of forces would necessarily imply a global balance of body velocities such that the wave regions would backslide as much as the interwave regions advance, and the worm would not translocate either forwards or backwards (47) (at least to first order in body strain *ε*, see mathematical analysis in (48)).

Mathematical modeling has shown that limbless locomotion with retrograde peristaltic waves as observed in our experiments can occur if substrate-anchoring is achieved at the wave regions (49). This theory is consistent with our measurements of *S. mansoni*, but it is unclear how the parasite is able to spatiotemporally coordinate muscular contraction with anchoring. The tegument (outer surface) of the adult schistosome parasite presents sexual dimorphism; it is covered in hemispherical tubercles each with actin spines in males, whereas it is relatively smooth in females (4,6,50). Given that adult males carry the females around the venous system, it has been suggested that the tubercles in the male worm tegument contribute to locomotion by anchoring the worm (6). The surface roughness elements provided by tubercles would locally increase the coefficient friction of the wave regions, where the worm body is thicker and presses against the substrate. The mechanism would be analogous to Coulomb friction, where the friction coefficient is proportional to the normal load acting on the contact surface. Numerical models of limbless crawling that include Coulomb-like substrate friction are able to recapitulate directional migration whereby the coordination between friction and contraction emerges via stick-slip transitions (51). Consistent with this idea, we have previously observed slip-stick transitions experimentally in *Physarum* fragments that crawl by generating peristaltic contraction waves. Furthermore, stick-slip friction has been proven mathematically to be more energy efficient than continuous linear friction in self-propelled crawlers (52).

### Locomotor similarities and differences between *S. mansoni* and other soft-bodied organisms

#### Sucker-based two-anchor crawlers

Crawling or marching mediated by two suckers can be found in other species *e.g*., leeches, that are challenged to remain attached to a surface (53). An interesting morphological difference between *S. mansoni* and the leech is that the posterior sucker is closely located to the anterior sucker in the schistosome, whereas, in the leech, the posterior sucker is located at the far end of the body. Thus, the leech has a longer stride relative to body length and a faster migration speed at the same stride frequency. However, having the two suckers at the front of the body is a more stable configuration when extending a soft extremity against flow. Conceivably, sucker-based support and locomotion are important for *S. mansoni* in the large-bore vessels of the hepatic portal system where the hemodynamic forces are the strongest that the parasite encounters.

Our measurements suggest that the suckers of *S. mansoni* can withstand a traction stress of ∼ 125 Pa before losing adherence. This value is in good agreement with previously reported measurements of maximum traction stresses for adherent single cells under flow (∼ 200 – 300 Pa, (34,54)), even though the mechanisms of substrate adhesion by *S. mansoni* and adherent cells are likely to be different. Our estimated detachment traction stress is, however, significantly lower than the adhesive pressure withstood by the much larger leech, *Whitmania pigra*, which was recently reported to be ∼50 KPa by Feng *et al* (55). The difference is explained by noting that, whereas Feng *et al* measured forces by pulling on the leeches in the direction perpendicular to the adhesion surface, we report force in the direction parallel to the surface, which reflects the capacity of the suckers to withstand shear rather than their generated suction pressure. We posit that the force in the direction parallel to the surface is more relevant for adhesion in *S. mansoni* given that the main hemodynamic force experienced by the worm is in the direction parallel to the vessel wall.

#### Peristaltic crawlers

Many terrestrial and marine worm genera, including the common earthworm, *Lumbricus terrestris*, use peristaltic waves to move over surfaces or burrow into the soil, and engage spine-like structures called setae to prevent the backsliding of the anchored body segments (56). Commonly, the setae are pointed posteriorly so that they provide higher resistance during backward motion than during forward motion. Numerical models have shown that in such cases, peristaltic waves of any gait lead to forward locomotion and there is no need for spatiotemporal coordination of contractility and friction (50,58,59). During penetration of the vertebrate host, schistosomula are covered with backward pointing spines (57), which likely provide the parasite with a robust mechanism for burrowing through the host’s skin. However, this apparent advantage comes at the price of preventing reversibility of locomotion by shifting the direction of wave propagation, as shown by mathematical analysis (49). By contrast, we have observed that adult male *S. mansoni*, which present isotropically-distributed tubercles on its surface, can spontaneously reverse its locomotion. It is reasonable to expect that the parasite exploits this mechanism to maneuver in narrow bore vessels where it would be impossible to reverse the direction of locomotion by turning its body around. In some species like *Lumbricus terrestris* and *Thoracophelia mucronata*, the hydroskeletal pressure rise caused by peristaltic contraction deploys setae only in the wave regions (20), thus achieving the desired spatiotemporal coordination between friction and contraction. Likewise, radial swelling in the peristaltic wave regions of adult *S. mansoni* could mediate expansion of the tubercles, thereby increasing the exposure of the spines to the substrate and the local friction coefficient. Both mechanisms would effectively create Coulomb-like friction which, as discussed above, may facilitate directional limbless crawling.

### Implications for drug discovery

Currently, treatment of schistosomiasis relies on just one drug, a rather insecure situation should the drug fail. The apparatus designed and approaches used here have identified and quantified a number of new biological parameters relating to the schistosome flatworm that could be employed to measure drug action, *e.g*., sucker grabbing strength and detachment rate under flow, body wave statistics, movement velocity, traction stress, and body-wave to sucker dynamics. As the mechanism of action of many anthelmintic drugs involves dysregulation of neuromuscular homeostasis (21) or, in the specific case of the schistosome parasite, disruption of the organization of the tegument including its component tubercles (58-60), the ability to measure these parameters under no-flow, flow and/or confinement conditions offers an unprecedented degree of sophistication. This new capacity could complement recent advances in the multi-parametric characterization of schistosome phenotypic space in response to drug insult (8,61-63). Future efforts shall focus on establishing proof-of-concept of these parameters for drug discovery with particular attention to their sensitivity and specificity, and address the scalability of the proposed methodologies to higher throughput.

## Conclusion

The present work identifies and measures the multi-modal ability of the mature schistosome parasite to undergo locomotion under various degrees of flow and confinement. Under no-flow and flow without confinement, the parasite employs its two suckers to move, whereas as confinement increases, this modality transitions to propulsion via peristaltic waves with/without the use of suckers. Our data suggests that locomotor mechanics might have been an important driver of the specific morphological features of the mature male schistosome, such as its tegument topography, as well as the strength and disposition of its suckers.

## Acknowledgements

We thank Brian Suzuki for expert assistance with the schistosome life-cycle. E.C.-H. would like to thank ‘la Caixa’ (Caja de Ahorros y Pensiones de Barcelona) for partial financial support through a ‘la Caixa’ Fellowship.

